# Willingness to Pay in the Human Brain: A fMRI Activation Likelihood Estimation Meta-analysis

**DOI:** 10.1101/2021.06.14.448018

**Authors:** Xiaoqiang Yao, Zhigang Huang, Yiwen Wang

## Abstract

The neural substrate of willingness to pay (WTP) ultimately supports human economic exchange activities and plays a crucial role in daily life. This paper aimed to identify the neural basis of WTP for food and nonfood, as well as the brain regions related to real and hypothetical WTP choices. We found the human brain centers of WTP by performing an activation likelihood estimation (ALE) meta-analysis (27 experiments, 796 subjects) on the existing neuroimaging studies. The conjunction analysis revealed that WTP for food and nonfood engaged a common cluster in the paracingulate and cingulate gyrus, revealing a common reward circuit in the brain. The frontal medial cortex and paracingulate gyrus were particularly activated by WTP for nonfood. Furthermore, the left caudate, left thalamus, angular gyrus and supramarginal gyrus (subregions of inferior parietal lobule) were more convergently activated by hypothetical WTP choice. Our findings support the idea that a common currency representation in the brain and reward-specific neural basis. Results also provide evidence of neural representations of the hypothetical bias.

## 1. Introduction

Consumers’ willingness to pay (WTP) plays a crucial role in daily life and is the cornerstone of marketing strategy such as developing competitive strategies, conducting to value audits, and designing optimal pricing for new product (Anderson, Jain, & Chintagunta, 1992). From an economic view, WTP denotes the maximum price a consumer calculated willing to pay for a given quantifies of a good (Schmidt & Bijmolt, 2020; Wertenbroch & Skiera, 2002). A WTP value reflects the consumer’s subjective value of the quantify of the good (Wertenbroch & Skiera, 2002). With the development of neuroimaging techniques and intersection of multiple disciplines, how the brain performs value computation has attracted the attention of scientist from decision and consumer neuroscience (Chib, Rangel, Shimojo, & O’Doherty, 2009; Clithero, Carter, & Huettel, 2009; De Martino, Kumaran, Holt, & Dolan, 2009; Hare, O’doherty, Camerer, Schultz, & Rangel, 2008; Plassmann, O’doherty, & Rangel, 2007).

What are our brain responses when we are making economic decisions? These studies revealed that activity in several areas including the medial orbitofrontal cortex (mOFC), anterior cingulate cortex (ACC), ventromedial prefrontal cortex (vmPFC), and inferior frontal gyrus (IFG), among others were correlated with economic decision-making (Chib et al., 2009; De Martino et al., 2009; Hare et al., 2008; Kennerley, Walton, Behrens, Buckley, & Rushworth, 2006; Padoa-Schioppa & Assad, 2006; Padoa-Schioppa & Conen, 2017; Wallis, 2007). Plassmann et al. (2007) firstly, investigated the neural representation of WTP in everyday economic transactions. They found that mOFC, ACC, and dorsolateral prefrontal cortex (DLPFC) were correlated with WTP. Likewise, Grueschow, Polania, Hare, and Ruff (2015) examined the neural representing of WTP for movie by using Becker-DeGroot-Marschak(BDM) auction (Becker, DeGroot, & Marschak, 1964) and they found that significant activity in the mPFC, the posterior cingulate cortex (PCC), and the parietal lobule among others during WTP decisions. However, the above studies have identified multiple brain regions associated with WTP computation, it is not clear how consistent and specific these findings are. In other words, whether there are specific brain regions associated with the computation of WTP, and the specificity of brain activity for different types of rewards and in distinct WTP measure contexts.

Previous studies on the neural mechanism of WTP have used different reward stimuli, which can be broadly classified as primary (i.e., food, sex) and secondary rewards (i.e., money, power). Primary rewards have innate value and are essential for human survival and reproduction, whereas secondary rewards are not directly related to survival (Sescousse, Caldú, Segura, & Dreher, 2013). Several studies have found that a phylogenetically and ontogenically older region posterior lateral OFC processes primary rewards for specificity, whereas a phylogenetically recent structure anterior lateral OFC processes secondary rewards for specificity (Sescousse et al., 2013; Sescousse, Redouté, & Dreher, 2010). There is also a common set of brain areas associated with processing primary and secondary rewards, including the ventral striatum, the anterior cingulate cortex, the anterior insula and among others (Sescousse et al., 2013; Sescousse et al., 2010). For example, Chib et al. (2009) found the vmPFC was correlated with subject’s WTP for all classes of items (money, trinkets and snacks). Other studies have also investigated the neural basis of WTP for non-primary rewards, such as movie and books. Grueschow et al. (2015) examined the neural representing of WTP for movie by using BDM auction and found that significant activity in the medial prefrontal cortex (mPFC), the posterior cingulate cortex (PCC), and the parietal lobule among others during WTP decisions. Likewise, Waskow et al. (2016) revealed that neural activity in the OFC, mPFC, and the ACC during decision-making correlates with individual’s WTP for music. Therefore, when performing WTP calculations for different reward types, are there common and different neural representations?

Another factor that affects the neural basis of WTP may be the measure context. There are various methods of measuring WTP in existing market research. According to the measure context, the methods of measuring WTP can be divided into hypothetical and real measuring of WTP (Miller, Hofstetter, Krohmer, & Zhang, 2011). Specifically, the BDM auction is the most widely used measures of WTP under real context (Becker et al., 1964). In the BDM procedure, participants first state their bids for each item, and then at the end of procedure, one bids will be randomly selected. If participants’ bid is less than the price of a randomly item, the participant does not buy the item, if participants’ bid is equal to or more than the price of a randomly item, the participant must buy the item for the randomly item price. The open questioning is one of the most used measure of WTP under hypothetical context, in which a participant stated their bids for each item. Real choice may usually be precise, direct, have greater stakes, and tend to be emotional, while hypothetical choices, without any consequences, may be quick and unconscious, and require fewer cognitive resources (Kang, Rangel, Camus, & Camerer, 2011).

Moreover, previous research has found a hypothetical bias between real and hypothetical WTP, that is, participants overstate their WTP for the goods under hypothetical conditions compared to in real WTP context (Schmidt & Bijmolt, 2020). They revealed that the hypothetical WTP is typically overestimated by almost 21% when compared to the real WTP for consumer goods (Schmidt & Bijmolt, 2020). Meanwhile, researchers also tested the neural representation mechanism of this hypothetical bias (Kang & Camerer, 2013; Kang et al., 2011). Kang et al. (2011) revealed that activity in OFC and ventral striatum (VS) correlated with the WTP for consumer products in both types of real and hypothetical decision. In addition, they found that the mOFC, ACC, caudate and inferior frontal gyrus were more responsive to decision value in real than hypothetical WTP. Conversely, Kang and Camerer (2013) found that more deactivation in the striatum and mOFC in real WTP for aversive foods, compared with hypothetical trials. Overall, whether there are brain regions activity specificity and consistency for real and hypothetical WTP have not been addressed. Answering these questions is important for understanding the neural basis of the WTP. To address the above issues, we supposed a quantitative meta-analysis of these studies by using different reward type and contexts of measuring WTP may be able to answer these questions above. Therefore, the aim of this study was to (1) examine the neural representation of WTP for different rewards, (2) to investigate the common and distinct of brain network of WTP for different rewards, and (3) to identify the consistency and specificity of WTP measures context neural basic in the brain. To achieve these aims, we conducted meta-analyses with activation likelihood estimation (ALE) method by comparing the WTP for food and nonfood reward as well as real and hypothetical WTP choices.

## 2. Materials and methods

### 2.1. literature search and selection

A systematic literature search was performed in Web of Science and PubMed databases to identify relevant studies (up to the August, 2020). The following key words were used for the search: “willingness to pay”, and “Becker-DeGroot-Marshak”, in combination(“And”) with “fMRI”. In addition to Web of science and PubMed search, several other sources were explored, including (a) work cited in review paper (Peters & Büchel, 2010); (b) cited articles in meta-analyses (Bartra, McGuire, & Kable, 2013; Levy & Glimcher, 2012); and (c) relevant reference lists of the initial search pre-selected articles.

The inclusion criteria for articles were as follows: (1) fMRI was used as the main methodology; (2) study that included at least one group of healthy participants were including; (3) fMRI results were derived from whole-brain analysis; (4) activations coordinates (foci) were presented in a standardized stereotaxic space (Talairach or Montreal Neurological Institute, MNI); (5) articles were published in peer-review journals; and (6) articles were written in English. (6) for each experiment per subject group, only one most strongly reflects the WTP process contrast’s coordinate data was extracted.

A total 25 studies (27 experiments) were included in following meta-analyses: 17 of these studies (19 experiments) focused on WTP for food reward, 11 of these studies (11 experiments) on WTP for nonfood reward (three studies contributed to both categories). On the other hand, according to the context of measurement WTP, it can be divided into hypothetical and real WTP. A real measure of WTP such as BDM auction, and a hypothetical measure such as open questioning (Miller et al., 2011). In this, there are 22 articles (24 experiments) that use the BDM method to measure real WTP, and 4 articles (4 experiments) that use the hypothetical method to measure WTP (one study contributed to both categories). Figure. 1 showed a Preferred Reporting Items for Systematic and Meta-Analyses (PRISMA) flowcharts depicts the steps taken to identification and identify eligible studies. Table 1 presents the details of all included studies. The present meta-analyses study was preregistered on the Open Science Framework (https://osf.io/6fbwr).

**Fig. 1.**
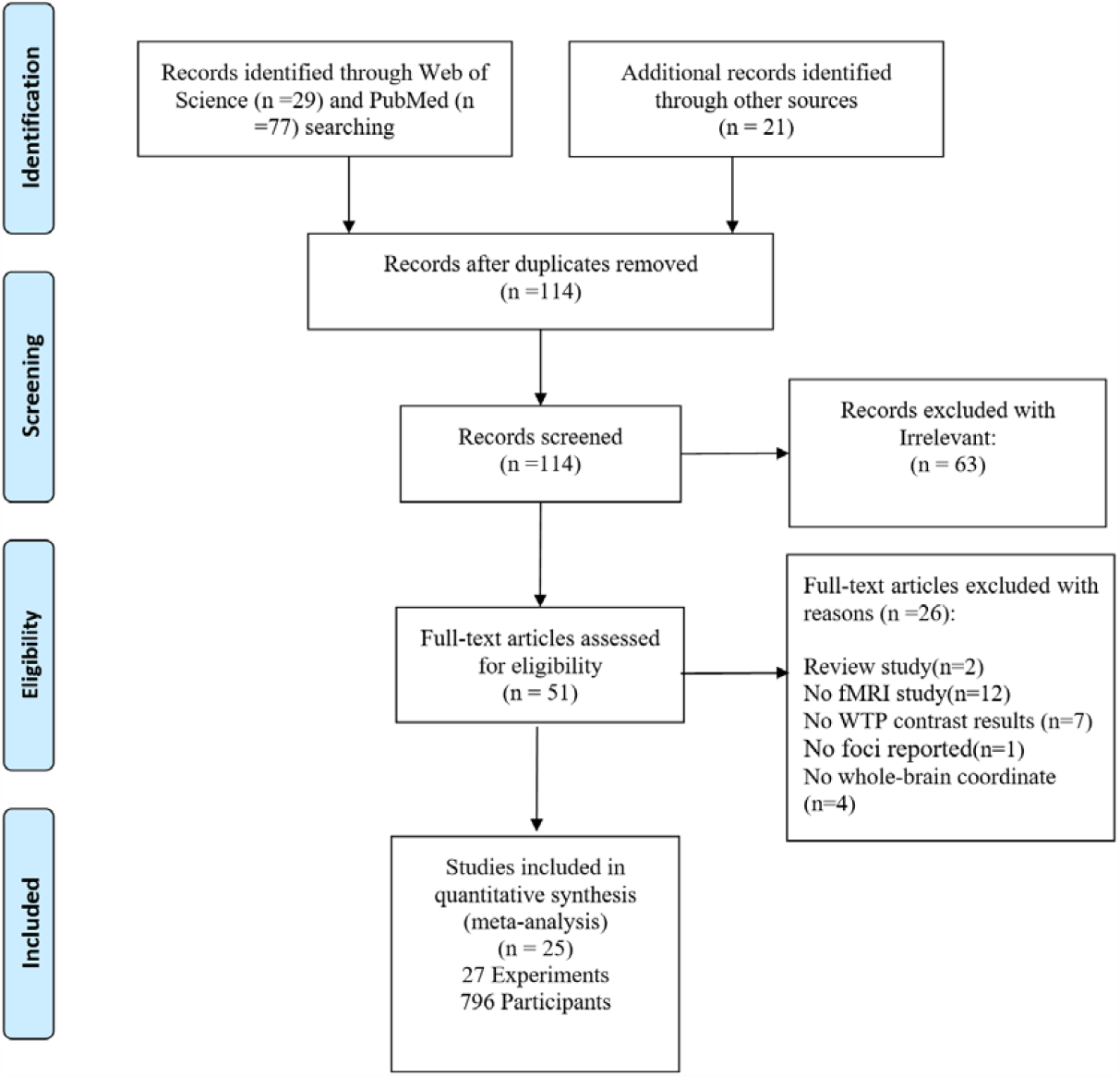
PRISMA flowchart of identification and identify eligible studies; fMRI, functional magnetic resonance imaging; WTP, willingness to pay.

**Table 1.** Overview of studies included for the meta-analysis.

| Article | Subjects(male) | Mean age | Methodology | Stimuli | Contrast included | Space | soft |
| --- | --- | --- | --- | --- | --- | --- | --- |
| Chib et al. (2009) S1 | 19(15M) | 23 | fMRI | Money/food/trinkets | Brain regions encode the WTP | MNI | SPM8 |
| Chib et al. (2009) S2 | 13(11M) | NA | fMRI | snacks | Activations for bid choices | MNI | SPM8 |
| Contreras-Rodriguez et al. (2020) | 75(NA) | 25-40 | fMRI | Foods | Functional > standard food trials | MNI | SPM |
| Enax et al. (2015a) | 33(NA) | NA | fMRI | Food products | Correlation with WTP | MNI | SPM8 |
| Enax et al. (2015b) | 25(11M) | 23.3 | fMRI | Food products | Correlates of WTP | MNI | SPM8 |
| Gospic et al. (2014) | 38(NA) | NA | fMRI | Charity picture | proposals > non-proposals | MNI | SPM8 |
| Grueschow et al. (2015) | 26(13M) | 20-28 | fMRI | DVD | Correlation with purchasing decisions | MNI | SPM8 |
| Hare et al. (2010) | 22(0M) | 24.7 | fMRI | Charitable organizations | Correlated with donation during free trials | MNI | SPM5 |
| Hare et al. (2008) | 16(9M) | 24.1 | fMRI | Foods | Activity in th goal values | MNI | SPM5 |
| Plassmann et al. (2007) | 19(16M) | 25.45 | fMRI | Foods | Correlates of WTP in free trials | MNI | SPM5 |
| Plassmann et al. (2010) S1 | 19(15M) | 23.7 | fMRI | Neutral and aversive foods | Free bid > force-bid trials | MNI | SPM5 |
| Plassmann et al. (2010) S2 | 20(15M) | 23-25 | fMRI | Aversive and appetitive foods | Correlation with free-bid trials | MNI | SPM5 |
| Lawn et al. (2020) | 38(29M) | NA | fMRI | Cigarettes | Purchase vs Not Purchase | MNI | SPM12 |
| Linder et al. (2010) | 30(15M) | 26.03 | fMRI | Foods | Correlated with WTP | MNI | SPM5 |
| Mackey et al. (2016) | 19(9M) | 31.5 | fMRI | Aversive and appetitive foods | WTP task-related regions | MNI | AFNI |
| De Martino et al. (2009) | 18(10M) | NA | fMRI | Lottery tickets | Correlated with WTP | MNI | SPM2 |
| Medic et al. (2014) | 47(23M) | 23.8 | fMRI | Snack items | Correlated with subjective value | MNI | SPM8 |
| Motoki et al. (2019) | 27(21M) | 20.37 | fMRI | Book | WTP during the purchasing task | MNI | SPM8 |
| Rihm et al. (2019) | 32(32M) | 26.13 | fMRI | Snack foods and trinket | Brain regions activation for all goods | MNI | SPM12 |
| Tang et al. (2014) | 29(NA) | NA | fMRI | Food | Correlated with participants bid | MNI | SPM8 |
| Verdejo-Román. et al. (2017) | 76(NA) | NA | fMRI | Food | Activations during WTP | MNI | SPM8 |
| Waskow et al. (2016) | 25(13) | 35.08 | fMRI | Music album | Correlated with the prices paid | MNI | SPM8 |
| Weber et al. (2007) | 16(10M) | 24.85 | fMRI | MP3 songs | Buying > Selling | MNI | SPM2 |
| Huijsmans et al. (2019) | 47(20M) | 23.5 | fMRI | Supermarket products | Abundance > scarcity | MNI | SPM12 |
| Winston et al. (2014) | 21(8M) | 24.0 | fMRI | Pain shocks | Correlation with bids | MNI | SPM8 |
| Zangemeister et al. (2019) | 22(NA) | NA | fMRI | Flavored liquid | Correlated with WTP | MNI | SPM8 |
| Kang et al. (2011) | 24(24M) | 20.9 | fMRI | Consumer products | Correlate with stimulus value | MNI | SPM5 |

**Note:** S1=study1; S2=study2; NA=Not available; fMRI=Functional magnetic resonance imaging; WTP= willingness to pay; MNI=Montreal Neurological Institute.

### 2.2. Activation likelihood estimation (ALE) method

#### 2.2.1. Activation likelihood estimation analyses

We performed the activation likelihood estimation (ALE) algorithm for coordinate-based meta-analysis (Eickhoff et al., 2009; Laird, Eickhoff, et al., 2009; Laird, Lancaster, & Fox, 2009; Turkeltaub, Eden, Jones, & Zeffiro, 2002), which aims to identify converging areas of reported coordinates across different experiments, empirically determining whether this clustering is higher than expected by chance (Eickhoff, Bzdok, Laird, Kurth, & Fox, 2012; Turkeltaub et al., 2012). The ALE algorithm to treat each focus reported on function neuroimaging studies as the center of a 3D Gaussian probability distribution and then calculates the maximum of each focus within an experiment, creating modeled activation maps. This ALE map is determined against the null hypothesis by using permutation tests (Eickhoff et al., 2012). For all analyses, the threshold for significance using a cluster-level family-wise error (cFWE) correction at *p*<0.05 with a cluster defining threshold of *p*<0.001 and 5,000 permutations (Eickhoff et al., 2012). To perform ALE meta-analysis, we used Ginger ALE v.3.0.2. (http://www.brainmap.org/ale/) with MNI coordinates.

#### 2.2.2. Conjunction and contrast analyses

To test neural basis that is common or distinct to WTP for food and nonfood rewards, and measure WTP under different context (hypothetical and real measure of WTP), we further performed conjunction (i.e., WTP for food and nonfood rewards; hypothetical and real WTP) and contrast analyses based on the ALE results (i.e., WTP for food reward > WTP for nonfood reward; hypothetical WTP > real WTP).

For conjunction analysis, we implemented by using the conservative minimum statistic (Nichols, Brett, Andersson, Wager, & Poline, 2005) to identify regions consistently recruited across conditions. For the contrast analysis, we firstly conducted separate ALE analyses for each condition and computing the voxel-wise difference between the ensuring ALE maps (Eickhoff et al., 2011). Afterwards, all experiments were pooled and randomly divided into two groups of same size as that in the two original sets of experiments, reflecting the contrasted ALE analyses (Langner, Leiberg, Hoffstaedter, & Eickhoff, 2018). The ALE scores of these two randomly assembled groups were calculated, and the difference between these ALE scores were recorded for each voxel of the brain. Repeating this process 25,000 times then yielded the null-distribution of differences in ALE values between the two conditions. The true difference in the ALE values was then tested against this voxel-wise null-distribution of label-exchangeability, and threshold at a probability of *p*> 95% for true differences.

#### 2.2.3. Data visualization

We used probabilistic cytoarchitectonic maps of human brain to label our resulting coordinates to anatomical structures, as implemented in SPM Anatomy Toolbox v.3 (Eickhoff, Heim, Zilles, & Amunts, 2006; Eickhoff et al., 2007; Eickhoff et al., 2005). For visualization purposes, BrainNet Viewer (Xia, Wang, & He, 2013) was implemented.

## 3. Results

### 3.1 WTP for food and nonfood ALE meta-analyses

#### 3.1.1 WTP for food

The results of ALE meta-analysis of WTP for food are presented in Table 2 and displayed visually in Fig. 2A. The ALE analysis revealed that the regions with significant activity in WTP for food were the paracingulate gyrus, cingulate gyrus, frontal medial cortex, frontal orbital cortex, insular cortex, frontal pole, frontal operculum cortex and inferior frontal gyrus.

**Table 2.** ALE meta-analyses result of WTP for food and nonfood.

| Cluster no | Macroanatomical location | MNI coordinate |  |  | Z score | Volume(mm^3) |
| --- | --- | --- | --- | --- | --- | --- |
|  |  | x | y | z |  |  |
| WTP for food |  |  |  |  |  |  |
| 1 | Paracingulate Gyrus/Cingulate Gyrus | -2 | 42 | -6 | 6.43 | 2952 |
|  | Cingulate Gyrus | 2 | 36 | 4 | 3.67 |  |
|  | Frontal Medial Cortex/Subcallosal Cortex | -4 | 32 | -18 | 3.48 |  |
| 2 | Frontal Orbital Cortex/Frontal Pole | 30 | 28 | -10 | 5.12 | 1760 |
|  | Insular Cortex/Frontal Orbital Cortex | 30 | 22 | -6 | 4.70 |  |
| 3 | Insular Cortex/Frontal Operculum Cortex | -36 | 22 | 0 | 4.27 | 1192 |
|  | Insular Cortex/ Frontal Orbital Cortex | -34 | 14 | -10 | 3.90 |  |
| 4 | Frontal Pole/Inferior Frontal Gyrus | 44 | 42 | 12 | 6.28 | 1056 |
| WTP for nonfood |  |  |  |  |  |  |
| 1 | Paracingulate Gyrus/Cingulate Gyrus | -2 | 42 | -6 | 6.29 | 2560 |
|  | Frontal Medial Cortex/Frontal Pole | -6 | 50 | -14 | 3.63 |  |
|  | Frontal Pole/ Frontal Medial Cortex | -4 | 56 | -8 | 3.51 |  |

*Note* : MNI=Montreal Neurological Institute. All peaks were assigned to the most probable brain area by using the SPM Anatomy toolbox V3.

**Fig. 2.**
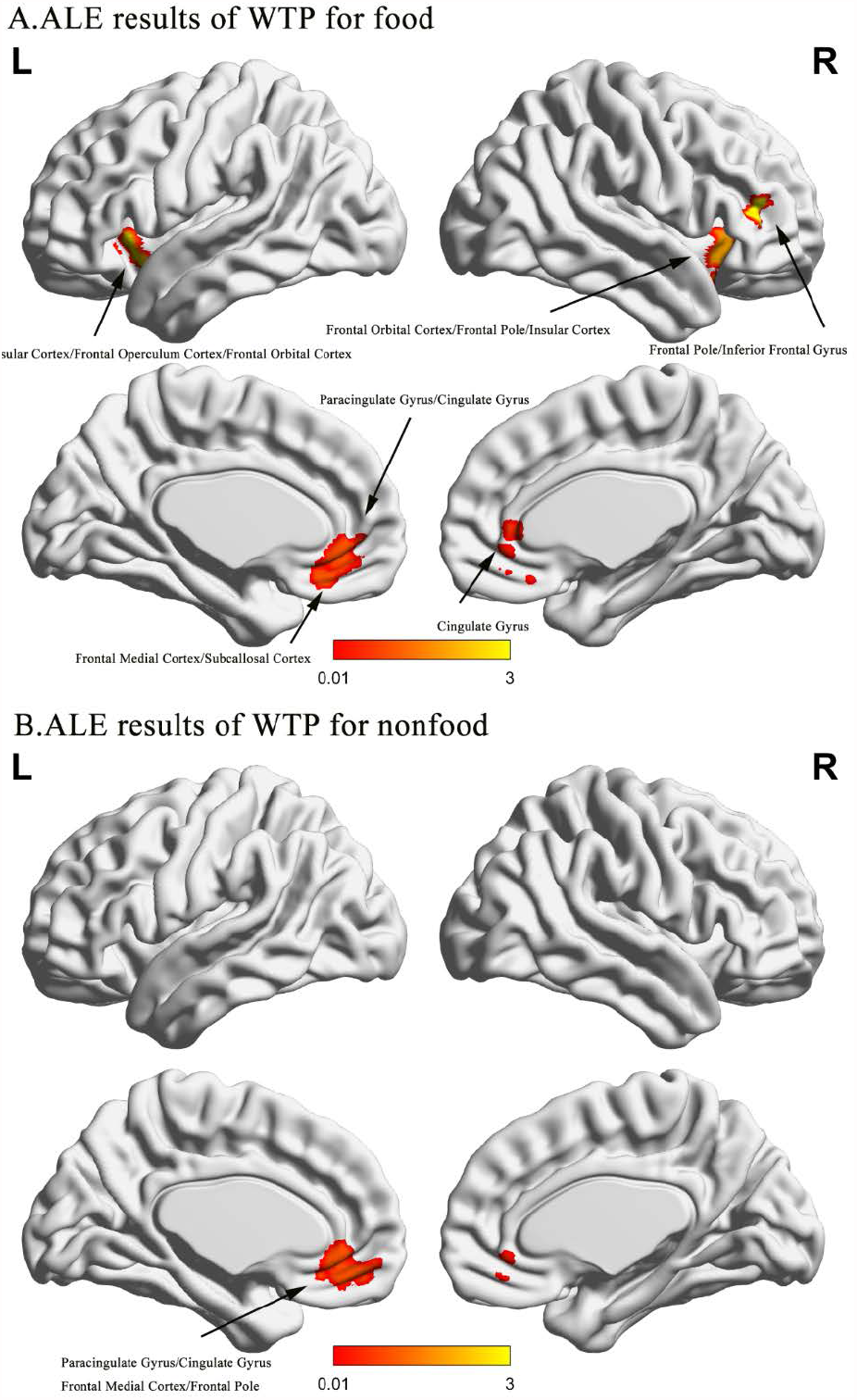
ALE results of WTP for food (A) and nonfood(B).

#### 2.1.2 WTP for nonfood

Results from ALE meta-analysis suggested that the regions with significant convergence activity in WTP for nonfood were the paracingulate gyrus, cingulate gyrus, frontal medial cortex, frontal pole, and frontal medial cortex (Table 2 and Figure 2B).

#### 2.1.3 Conjunction and contrast analyses of food and nonfood WTP

The results of conjunction and contrast with food and nonfood WTP are presented in Table 3 and displayed visually in Fig. 3. The conjunction analysis revealed that a common area was activated in the paracingulate gyrus and cingulate gyrus for both food and nonfood WTP (Fig. 3A). The contrast analysis revealed that the frontal medial cortex and paracingulate gyrus were stronger activity in nonfood WTP (Fig. 3B). However, the contrast analysis showed that no clusters was found for food WTP than nonfood WTP.

**Table 3.** Contrast and conjunction analyses of food and nonfood WTP.

| Cluster no | Macroanatomical location | MNI coordinate |  |  | Z score | Volume(mm^3) |
| --- | --- | --- | --- | --- | --- | --- |
|  |  | x | y | z |  |  |
| Food WTP ∩ nonfood WTP |  |  |  |  |  |  |
| 1 | Paracingulate Gyrus/ Cingulate Gyrus | -2 | 42 | -6 |  | 1824 |
| WTP for Food >nonfood |  |  |  |  |  |  |
| No clusters found |  |  |  |  |  |  |
| Nonfood WTP >Food WTP |  |  |  |  |  |  |
| 1 | Frontal Medial Cortex/ Paracingulate Gyrus | -6 | 50 | -16 | 2.61 | 208 |

*Note:* MNI=Montreal Neurological Institute. All peaks were assigned to the most probable brain area by using the
SPM Anatomy toolbox V3.

**Fig. 3.**
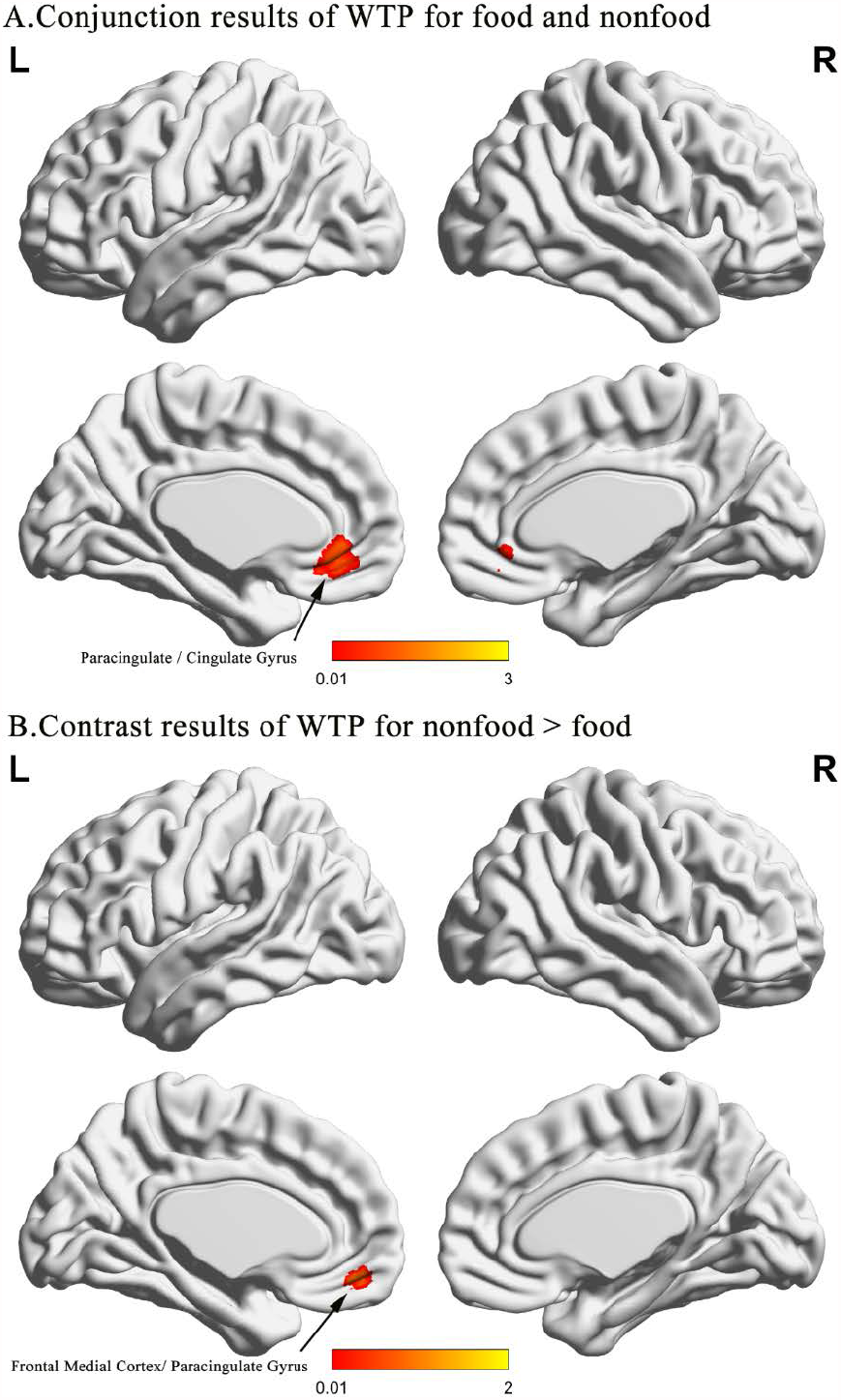
Results of the ALE conjunction (A) and contrast (B) analyses of food and nonfood WTP.

### 2.2 Real and hypothetical WTP choice ALE meta-analyses

#### 2.2.1 Real WTP choice

The regions with significant activations in real WTP choice are shown in Table 4 and Fig 4 A. Significant cross-study convergence activity during real WTP choice were observed in the bilateral cingulate gyrus, paracingulate gyrus, frontal pole, inferior frontal gyrus, insular cortex, frontal operculum cortex and frontal orbital cortex.

**Table 4.** ALE meta-analyses result of real WTP and hypothetical WTP choice.

| Cluster no | Macroanatomical location | MNI coordinate |  |  | Z score | Volume(mm^3) |
| --- | --- | --- | --- | --- | --- | --- |
|  |  | x | y | z |  |  |
| Real WTP choice |  |  |  |  |  |  |
| 1 | Paracingulate Gyrus/Cingulate Gyrus | -4 | 40 | -6 | 6.83 | 3760 |
|  | Cingulate Gyrus | 2 | 36 | 4 | 3.84 |  |
| 2 | Frontal Pole/Inferior Frontal Gyrus | 44 | 42 | 12 | 6.37 | 1096 |
| 3 | Frontal Orbital Cortex/Insular Cortex | 28 | 26 | -8 | 4.21 | 1016 |
| 4 | Insular Cortex / Frontal Operculum Cortex | -36 | 22 | 0 | 4.33 | 936 |
|  | Insular Cortex/ Frontal Orbital Cortex | -30 | 14 | -8 | 3.59 |  |
| Hypothetical WTP choice |  |  |  |  |  |  |
| 1 | Left Caudate/Left Accumbens | -6 | 4 | -2 | 5.13 | 528 |
| 2 | Angular Gyrus/Supramarginal Gyrus | 62 | -46 | 26 | 4.60 | 496 |

*Note* : MNI=Montreal Neurological Institute. All peaks were assigned to the most probable brain area by using the
SPM Anatomy toolbox V3.

**Fig. 4.**
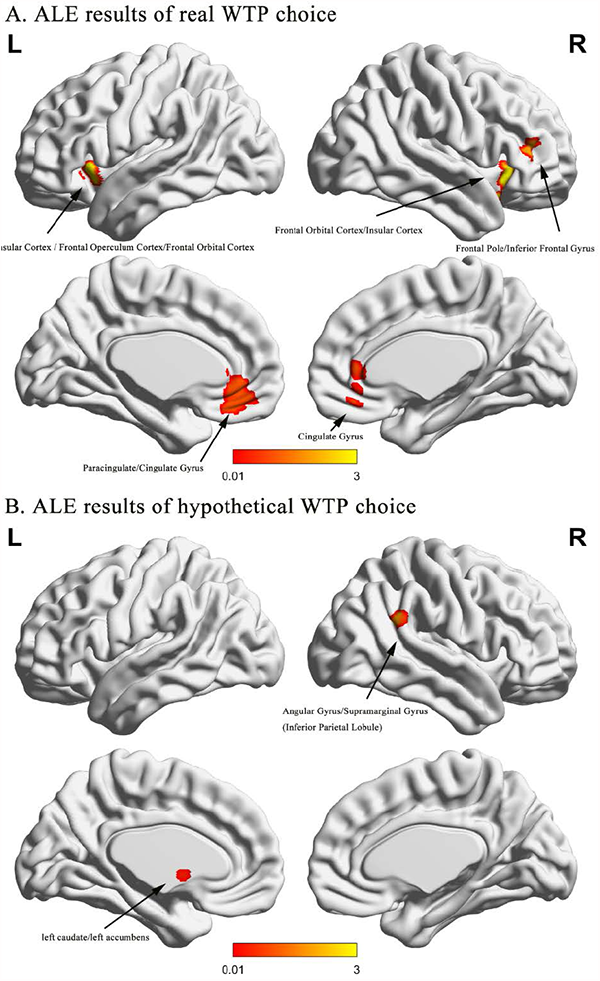
ALE results of real WTP (A) and hypothetical WTP choice (B).

#### 2.2.2 Hypothetical WTP choice

The regions with significant activations in hypothetical WTP choice are shown in Table 4 and Fig. 4B. The brain areas of significant cross-study activity were observed in the left caudate, left accumbens, angular gyrus (AG) and supramarginal gyrus (SMG), both AG and SMG are subregions of inferior parietal lobule (IPL) and overlap with temporoparietal junction (TPJ).

#### 2.2.3 Conjunction and contrast analyses of real and hypothetical WTP choices

The results of contrast and conjunction of real and hypothetical WTP choices are illustrated in Table 5 and Fig. 5. The results of conjunction analysis showed that no clusters found. The contrast analysis revealed that the left caudate, left thalamus, angular gyrus, and supermarginal gyrus (subregions of IPL) were stronger activity in hypothetical WTP choice (Fig. 5). However, the contrast analysis showed that no cluster found for real WTP than hypothetical WTP.

**Table 5.**
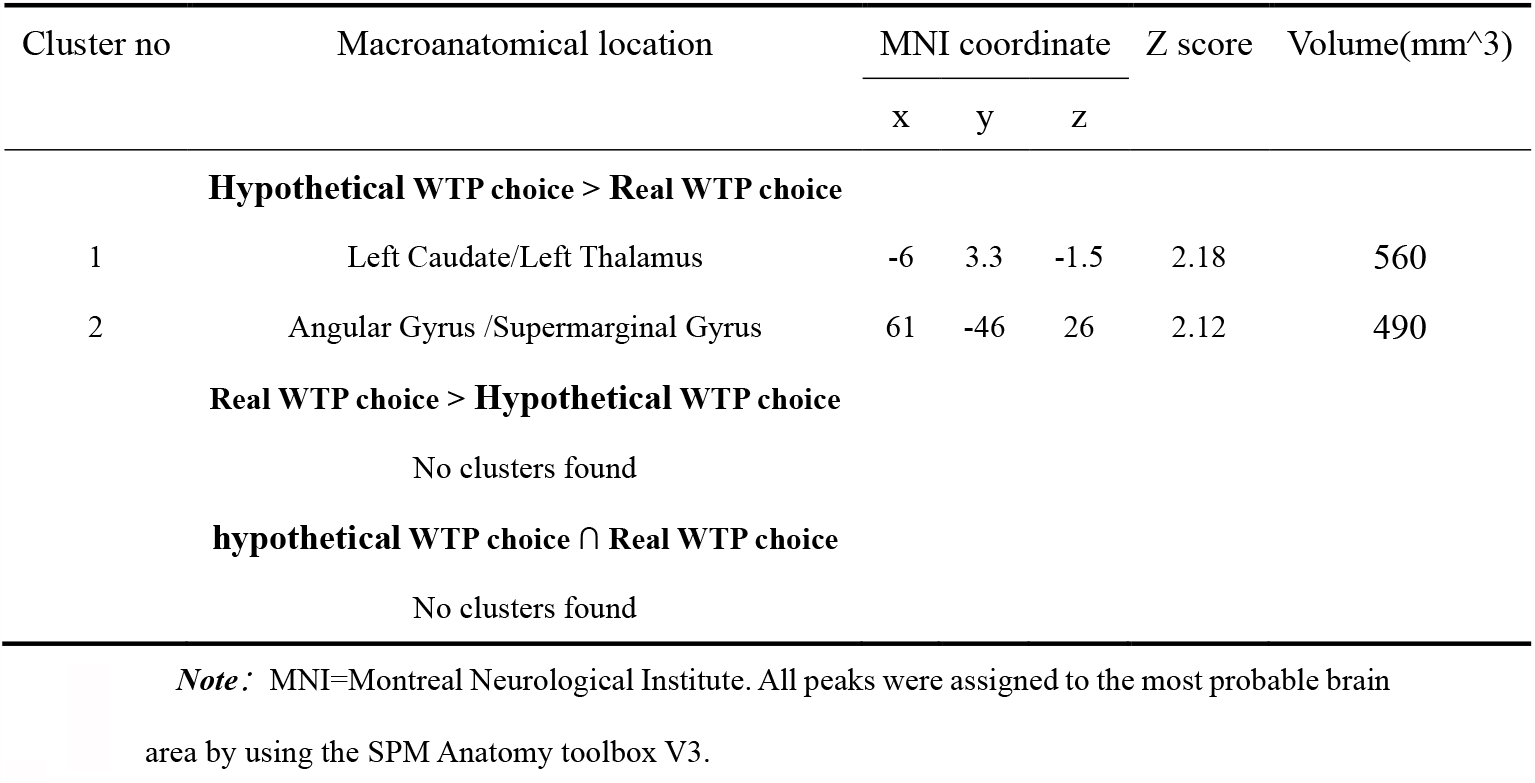
Contrast and conjunction analyses of real and hypothetical WTP choice.

**Fig. 5.**
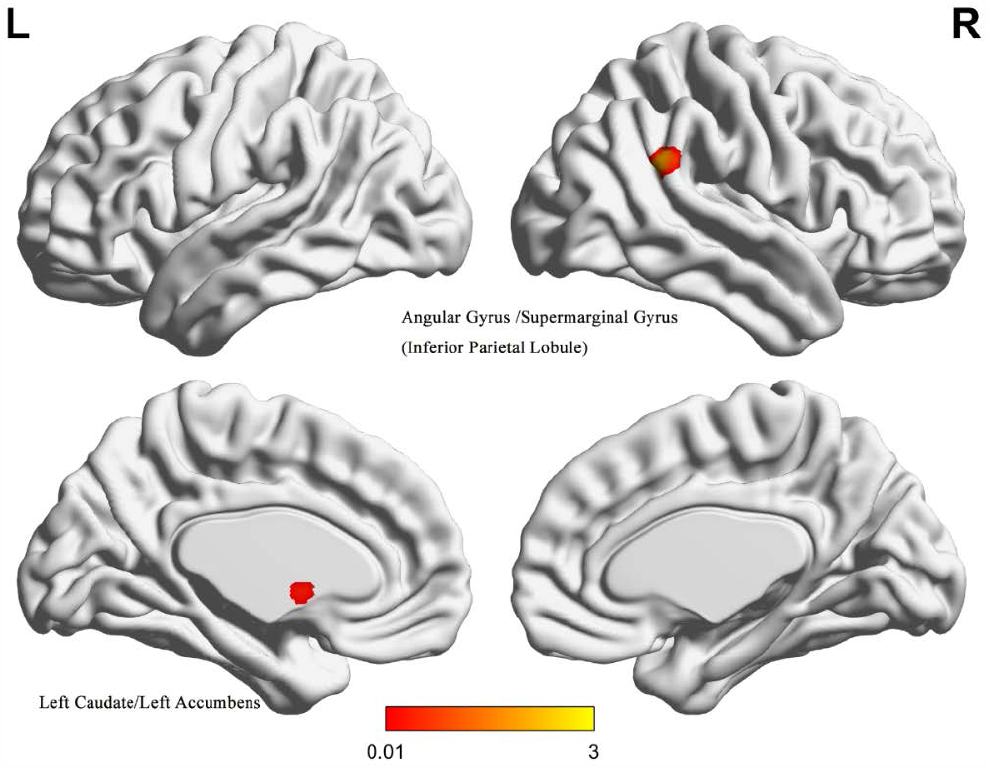
The results of the contrast analysis between the real and hypothetical WTP choice.

## 4. Discussion

In the present study we investigate the neural representation of WTP, as well as the specificity and consistency of brain activity across different types of reward and WTP measurement contexts. The results indicate there is a neural circuit consisting of the paracingulate gyrus, cingulate gyrus, frontal medial cortex, frontal orbital cortex, insular cortex, frontal pole, inferior frontal gyrus and among others significant activation during WTP for food and nonfood. ALE results demonstrated that the paracingulate and cingulate gyrus link to both food and nonfood reward WTP processing, reflecting a common neural representation of WTP. In contrast, the WTP for nonfood were more engaged in the frontal medial cortex and paracingulate gyrus. Moreover, real WTP choice engaged a neural network consisting of the paracingulate gyrus, cingulate gyrus, frontal pole, inferior frontal gyrus, frontal orbital cortex, and insular cortex. In addition, the hypothetical WTP choice specific responses were further observed in the left caudate, left thalamus, angular gyrus and supramarginal gyrus (subregions of IPL).

To be more specific, we first investigated the neural basis of WTP computation for different rewards, and identified the common and distinct brain regions of WTP for different rewards. Results showed that frontal medial cortex, frontal pole and cingulate gyrus and paracingulate gyrus were associated with WTP computation for both food and nonfood rewards. This result is similar to several previous studies on the neural mechanisms of decision value in response to different rewards (Chib et al., 2009; Kang et al., 2011; Rihm et al., 2019). These findings support the idea that a set of regions in the brain process of reward computation in an indiscriminate manner. This results also in line with the common currency hypothesis that frontal medial cortex encodes different types of decision values. Previous studies have highlighted the role of ACC in cognitive processes, including cost-benefit decision-making, motivation, and action-reward (Holroyd & Yeung, 2012; Rushworth, Noonan, Boorman, Walton, & Behrens, 2011; Shenhav, Botvinick, & Cohen, 2013). In addition, our results revealed that nonprimary reward specific activations in the frontal medial cortex and paracingulate gyrus. A similar result was found in a previous neuroimaging meta-analysis of primary and secondary rewards. Specially, Sescousse et al. (2013) found that anterior lateral OFC was more associated with processing secondary rewards and the OFC of the posterior lateral was more associated with processing primary rewards. The results of the current study may also support the idea that non-primary rewards processing is in the anterior lateral frontal lobe compared to primary reward. Lastly, inconsistent with the two previous meta-analyses (Bartra et al., 2013; Sescousse et al., 2013), VS was not found to be activated in the food and nonfood reward WTP calculations in the current study. Previous study reported that VS encode prediction errors (Hare et al., 2008). One possible explanation is that the absence of VS activation is due to the absence of feedback from WTP decisions in the studies included in the current meta-analysis, and therefore would not have included prediction errors and correct responses to WTP decision outcomes.

We also examined the neural representations of both real and hypothetical WTP choice. Our contrast analysis suggested that hypothetical WTP choice specific responses in the left caudate, left thalamus, angular gyrus and supramarginal gyrus (subregions of IPL). This result is consistent with multiple previous studies of WTP for goods measured using hypothetical methods (Contreras-Rodriguez et al., 2020; Kang et al., 2011; Verdejo-Román, Fornito, Soriano-Mas, Vilar-López, & Verdejo-García, 2017). Verdejo-Román et al. (2017) found that compared with plain food processing of highly palatable food was associated with greater activation of supramarginal gyrus. Another study by Contreras-Rodriguez et al. (2020) revealed that angular gyrus increased activation during WTP for functional food compared with standard food. Lastly, Kang et al. (2011) found stronger activation in processing for hypothetical than real WTP choice in left caudate. This result is similar to the findings by Kang and Camerer (2013), which found stronger activation in some brain regions in the hypothetical WTP choice compared to the WTP real choice, such as left thalamus. Therefore, we hypothesized that these regions could be reflecting the brain representation of “hypothetical bias” in the WTP computation.

Besides, some limitations of this study need to be discussed as well. First, the current study has relatively fewer foci in the hypothetical WTP choice compared to the foci in the real WTP choice. Moreover, the different types of rewards in the real and hypothetical WTP conditions which may also affect the robustness of the results. This makes us need to be more cautious when interpreting and generalizing this result. Second, the current meta-analysis classified the types of stimuli used in the included studies into primary and non-primary rewards. Although these studies that have used food stimuli as reward, there was some heterogeneity among them. For example, some studies selected rewards for appetite foods and one for aversive foods. Therefore, it is uncertain whether there are different neural representations between these specific stimuli.

Taken together, our results demonstrated that a set of brain region was correlate with WTP for food and nonfood rewards computation. These results confirm that a widespread neural circuit contributes to the WTP process. Moreover, the results revealed the distinct and common neural basic on the WTP for different rewards. Lastly, our results provide evidence from neural representations for the hypothetical bias.

## Footnotes

1 Asterisk denotes articles included in the meta-analysis.

